# Sequence Specific Modeling of *E. coli* Cell-Free Protein Synthesis

**DOI:** 10.1101/139774

**Authors:** Michael Vilkhovoy, Nicholas Horvath, Che-Hsiao Shih, Joseph A. Wayman, Kara Calhoun, James Swartz, Jeffrey D. Varner

**Affiliations:** Robert Frederick Smith School of Chemical and Biomolecular Engineering, Cornell University, Ithaca, NY 14853; Davidson School of Chemical Engineering, Purdue University, West Lafayette, IN 47907; School of Chemical Engineering, Stanford University, Stanford, CA 94305

**Keywords:** Synthetic biology, constraint based modeling, cell-free protein synthesis

## Abstract

Cell-free protein synthesis (CFPS) is a widely used research tool in systems and synthetic biology. However, if CFPS is to become a mainstream technology for applications such as point of care manufacturing, we must understand the performance limits and costs of these systems. Toward this question, we used sequence specific constraint based modeling to evaluate the performance of *E. coli* cell-free protein synthesis. A core *E. coli* metabolic network, describing glycolysis, the pentose phosphate pathway, energy metabolism, amino acid biosynthesis and degradation was augmented with sequence specific descriptions of transcription and translation and effective models of promoter function. Model parameters were largely taken from literature, thus the constraint based approach coupled the transcription and translation of the protein product, and the regulation of gene expression, with the availability of metabolic resources using only a limited number of adjustable model parameters. We tested this approach by simulating the expression of two model proteins: chloramphenicol acetyltransferase and dual emission green fluorescent protein, for which we have training data sets; we then expanded the simulations to a range of additional proteins. Protein expression simulations were consistent with measurements for a variety of cases. The constraint based simulations confirmed that oxidative phosphorylation was active in the CAT cell-free extract, as without it there was no feasible solution within the experimental constraints of the system. We then compared the metabolism of theoretically optimal and experimentally constrained CFPS reactions, and developed parameter free correlations which could be used to estimate productivity as a function of protein length and promoter type. Lastly, global sensitivity analysis identified the key metabolic processes that controlled CFPS productivity and energy efficiency. In summary, sequence specific constraint based modeling of CFPS offered a novel means to *a priori* estimate the performance of a cell-free system, using only a limited number of adjustable parameters. While we modeled the production of a single protein in this study, the approach could easily be extended to multi-protein synthetic circuits, RNA circuits or the cell free production of small molecule products.

## 1 Introduction

Cell-free protein expression has become a widely used research tool in systems and synthetic biology, and a promising technology for personalized protein production. Cell-free systems offer many advantages for the study, manipulation and modeling of metabolism compared to *in vivo* processes. Central amongst these is direct access to metabolites and the biosynthetic machinery without the interference of a cell wall or the complications associated with cell growth. This allows interrogation of the chemical environment while the biosynthetic machinery is operating, potentially at a fine time resolution. Cell-free protein synthesis (CFPS) systems are arguably the most prominent examples of cell-free systems used today (*1*). However, CFPS is not new; Matthaei and Nirenberg first used E. *coli* cell-free extracts in the 1960s to decipher the sequencing of the genetic code (*2, 3*). Spirin and coworkers later improved the operational lifetime of cell-free protein production with a continuous exchange of reactants and products; however, these systems could only synthesize a single product and were energy limited (*4*). More recently, CFPS was improved by generating ATP using both substrate level (*5*) and oxidative phosphorylation (*6, 7*). Today, cell-free systems are used in a variety of applications ranging from therapeutic protein production (*8, 9*) to synthetic biology (*10*). There are also several CFPS technology platforms, such as the PANOx-SP and Cytomin platforms developed by Swartz and coworkers (*1, 5, 6*), and the TX/TL platform of Noireaux (*11*). However, if CFPS is to become a mainstream technology for advanced applications such as point of care manufacturing (*12*), we must first understand the performance limits and costs of these systems (*1*). One tool to address these questions is constraint based modeling.

Constraint based approaches such as flux balance analysis (FBA), which use stoichiometric reconstructions of microbial metabolism, have become standard tools in systems biology and metabolic engineering (*13*). FBA and metabolic flux analysis (MFA) (*14*), as well as convex network decomposition approaches such as elementary modes (*15*) and extreme pathways (*16*), model intracellular metabolism using the biochemical stoichiometry and other constraints such as thermodynamical feasibility (*17, 18*) under pseudo steady state conditions. Constraint based approaches have used linear programming (*19*) to predict productivity (*20, 21*), yield (*20*), mutant behavior (*22*), and growth phenotypes (*23*) for biochemical networks of varying complexity, including genome scale networks, using a limited number of adjustable parameters. Since the first genome scale stoichiometric model of *E. coli* (*24*), stoichiometric reconstructions of hundreds of organisms, including industrially important prokaryotes such as *E. coli* (*25*) and *B. subtilis* (*26*), are now available (*27*). Stoichiometric reconstructions have been expanded to include the integration of metabolism with detailed descriptions of gene expression (ME-Model) (*23, 28, 29*) and protein structures (GEM-PRO) (*30, 31*). These expansions have greatly increased the scope of questions that constraint based models can explore. Thus, constraint based methods are powerful tools to estimate the performance of metabolic networks. However, constraint based methods are typically used to model *in vivo* processes, and have not yet been applied to cell-free metabolism.

In this study, we used sequence specific constraint based modeling to evaluate the performance of *E. coli* cell-free protein synthesis. A core *E. coli* cell-free metabolic model describing glycolysis, pentose phosphate pathway, energy metabolism, amino acid biosynthesis and degradation was developed from literature (*25*); this model was then augmented with sequence specific descriptions of promoter function, transcription and translation processes. Thus, the sequence specific constraint based approach explicitly coupled transcription and translation processes with the availability of metabolic resources in the CFPS reaction. We tested this approach by simulating the cell-free production of two model proteins, and then investigated the productivity and energy efficiency for eight additional proteins. Productivity was inversely proportional to carbon number, while energy efficiency was independent of protein size. Based on these simulations, effective correlations for the productivity and energy efficiency as a function of protein length were developed. These correlations were then independently validated with a protein not in the original data set. Further, global sensitivity analysis identified the key metabolic processes that controlled CFPS performance; oxidative phosphorylation was vital to energy efficiency, while the translation rate was the most important factor controlling productivity. Lastly, we compared theoretically optimal metabolic flux distributions with experimentally constrained flux distributions; CFPS retained an *in vivo* operational memory that led to the overconsumption of glucose which negatively influenced energy efficiency. Taken together, sequence specific constraint based modeling of CFPS offered a novel means to *a priori* estimate the performance of a cell-free system, using only a limited number of adjustable parameters. While we considered only a single protein here, this approach could be extended to synthetic circuits, RNA circuits (*32*) or even cell-free small molecule production.

## 2 Results and discussion

### 2.1 Model derivation and validation

The cell-free stoichiometric network was constructed by removing growth associated reactions from the iAF1260 reconstruction of K-12 MG1655 E. *coli* (*25*), and adding deletions associated with the specific cell-free system (see Materials and Methods). The iAF1260 reconstruction describes 1260 ORFs, and thermodynamically derived metabolic flux directionality. We then added the transcription and translation template reactions of Allen and Palsson for the specific proteins of interest (*28*). A schematic of the metabolic network, consisting of 264 reactions and 146 species, is shown in Fig. 1A. The network described the major carbon and energy pathways and amino acid biosynthesis and degradation pathways. Using this network in combination with effective promoter models taken from Moon et al. (*33*) and literature values for cell-free culture parameters (Table 2), we simulated the sequence specific production of two model proteins: chloramphenicol acetyltransferase (CAT) and dual emission green fluorescent protein (deGFP, shown in the Supporting Information). We calculated the transcription rate using effective promoter models, and then maximized the rate of translation within biologically realistic bounds. Transcription and translation rates were subject to resource constraints encoded by the metabolic network, and transcription and translation model parameters were largely derived from literature (Table 2). In this study, we did not explicitly consider protein folding. However, the addition of chaperone or other protein maturation steps could easily be accommodated within the approach by updating the template reactions, see Palsson and coworkers (*23*). The cell-free metabolic model code and parameters can be downloaded under an MIT software license from the Varnerlab website (*34*).

**Figure 1:**
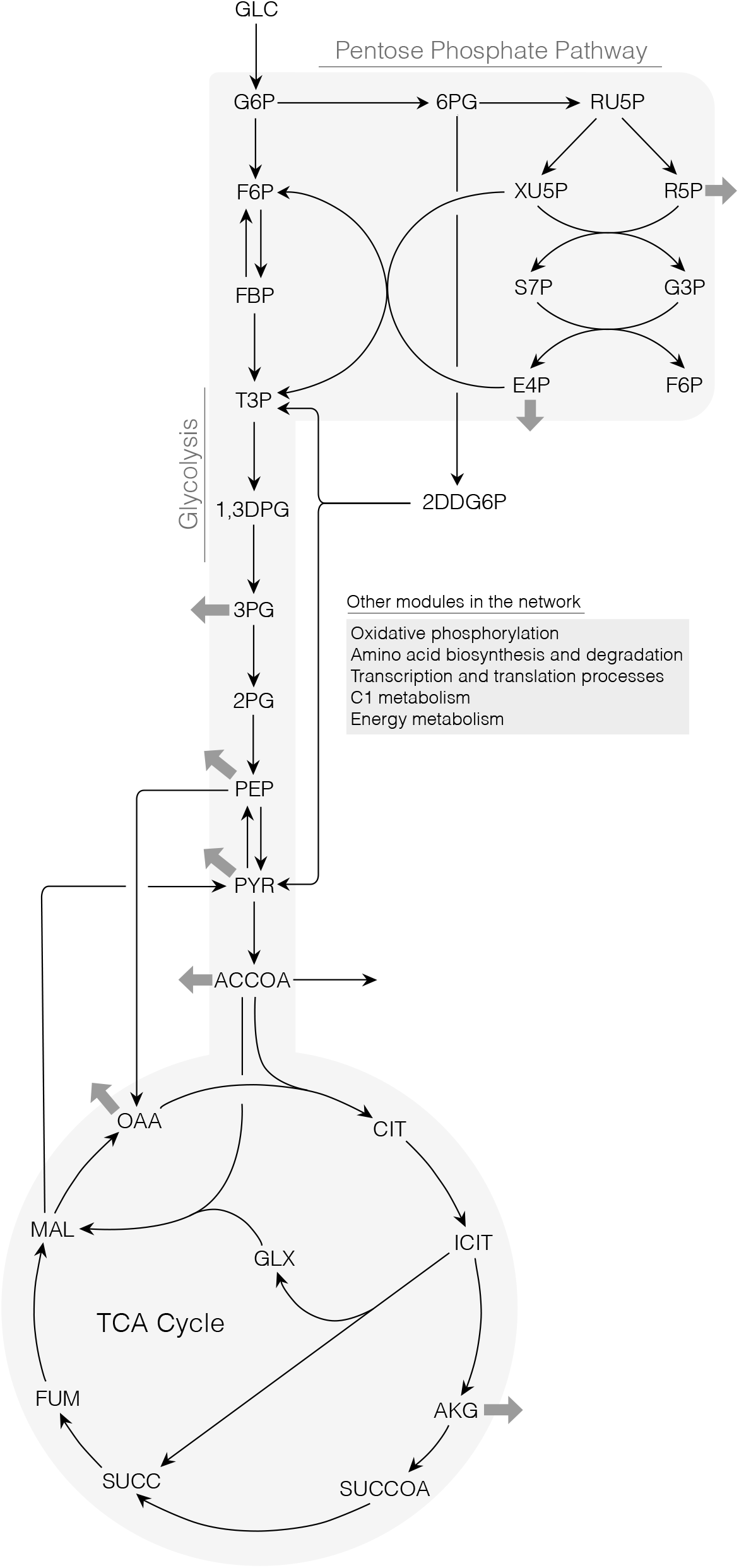
Sequence specific flux balance analysis. A. Schematic of the core metabolic network describing glycolysis, pentose phosphate pathway, the TCA cycle and the Entner-Doudoroff pathway. Thick gray arrows indicate withdrawal of precursors for amino acid synthesis.

**Table 1:** Transcription and translation template reactions for protein production. The symbol 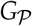 denotes the gene encoding protein product 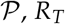, *R_T_* denotes the concentration of RNA polymerase, 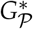 denotes the gene bounded by the RNA polymerase (open complex), *η_i_* and *α_j_* denote the stoichiometric coefficients for nucleotide and amino acid, respectively, *Pi* denotes inorganic phosphate, *R_X_* denotes the ribosome concentration, 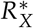 denotes bound ribosome, and *AA_j_* denotes *j^th^* amino acid.

| Description | Template reaction |
| --- | --- |
| Transcription initiation | $G_{\mathcal{P}} + R_T \longrightarrow G_{\mathcal{P}}^*$ |
| Transcription ($w_T$) | $G_{\mathcal{P}}^* + \sum_{k \in \{A,C,G,U\}} \eta_k \cdot (\{k\} TP + H_2O) \longrightarrow mRNA + G_{\mathcal{P}} + R_T + \sum_{k \in \{A,C,G,U\}} \eta_k \cdot PPi$ |
| mRNA degradation | $mRNA \longrightarrow \sum_{k \in \{A,C,G,U\}} \eta_k \cdot \{k\} MP$ |
| Translation initiation | $mRNA + R_X \longrightarrow R_X^*$ |
| tRNA charging | $\alpha_j \cdot (AA_j + tRNA + ATP + H_2O) \longrightarrow \alpha_j \cdot (AA_j-tRNA_j + AMP + PPi)$<br>$j = 1, 2, \dots, 20$ |
| Translation ($w_X$) | $R_X^* + \sum_{j \in \{AA\}} \alpha_j \cdot (AA_j-tRNA_j + 2GTP + 2H_2O) \longrightarrow \mathcal{P} + R_X + mRNA$<br>$+ \sum_{j \in \{AA\}} \alpha_j \cdot (tRNA + 2GDP + 2Pi)$ |

**Table 2:** Parameters for sequence specific flux balance analysis

| Description | Parameter | Value | Units | Reference |
| --- | --- | --- | --- | --- |
| T7 RNA polymerase concentration | $R_T$ | 1.0 | $\mu\text{M}$ | specified |
| Native RNA polymerase concentration | $R_T$ | 75 | nM | (11) |
| Ribosome concentration | $R_X$ | 1.6 | $\mu\text{M}$ | (11, 37) |
| Transcription elongation rate | $\dot{v}_T$ | 25 | nt/s | (11) |
| Translation elongation rate | $\dot{v}_X$ | 2 | aa/s/ribosome | (11, 37) |
| T7 transcription saturation coefficient | $K_{T7,T}$ | 116 | nM | estimated |
| P70 transcription saturation coefficient | $K_{P70,T}$ | 3.5 | nM | estimated |
| Translation saturation coefficient | $K_X$ | 45.0 | $\mu\text{M}$ | estimated |
| Polysome number | $K_P$ | 10 | ribosome number | estimated |
| mRNA degradation rate constant | $\lambda$ | 5.2 | $\text{h}^{-1}$ | (11) |
| T7 promoter weight | $K_{T7}$ | 10 | constant | estimated |
| Weight RNA polymerase binding alone P70a | $K_1$ | 0.014 | constant | estimated |
| Weight bound RNAP-$\sigma_{70}$ P70a | $K_2$ | 10 | constant | estimated |
| $\sigma_{70}$ concentration | $\sigma_{70}$ | 35 | nM | (11) |
| $\sigma_{70}$ dissociation constant | $K_D$ | 130 | nM | (54) |
| $\sigma_{70}$ hill coefficient | $n$ | 1 | constant | (54) |
| Gene concentration | $G_P$ | 5 | nM | (11) |
| ATP transcription coefficient (CAT) | $\text{ATP}_T$ | 176 | constant | calculated |
| CTP transcription coefficient (CAT) | $\text{CTP}_T$ | 144 | constant | calculated |
| GTP transcription coefficient (CAT) | $\text{GTP}_T$ | 151 | constant | calculated |
| UTP transcription coefficient (CAT) | $\text{UTP}_T$ | 189 | constant | calculated |
| ATP tRNA charging coefficient (CAT) | $\text{ATP}_X$ | 219 | constant | calculated |
| GTP translation coefficient (CAT) | $\text{GTP}_X$ | 438 | constant | calculated |

Cell-free simulations of the time evolution of CAT production were consistent with experimental measurements (Fig. 2). CAT was produced under a T7 promoter in a glucose/NMP cell-free system using glucose as a source of carbon and energy (*35*). Metabolic fluxes were constrained by experimental measurements of glucose, nucleotides, amino and organic acid consumption and production rates (estimated from a total of 37 metabolite time series measurements) for the first hour of the reaction (rates assumed constant; see Supporting Information). On the other hand, the rates of CAT transcription and translation were predicted by the model. The model showed good agreement with the CAT measurement with a coefficient of determination of R^2^ = 0.92. Next, we also simulated the production of deGFP under a P70a promoter in TXTL 2.0 using maltose and 3-phosphoglycerate (3PG) as a carbon and energy source (R^2^ = 0.84). The model captured the saturation of the deGFP titer for a range of plasmid concentrations (R^2^ = 0.97, see Supporting Information). Uncertainty in experimental factors such as the concentration of RNA polymerase, ribosomes, transcription and translation elongation rates, as well as the upper bounds on oxygen and carbon consumption rates (uniformly sampled around the parameter values shown in Table 2), did not qualitatively alter the performance of the model for both proteins (blue region, 95% confidence estimate). Together, these simulations suggested the description of transcription and translation, and its integration with metabolism encoded in the cell-free model, were consistent with experimental measurements. These simulations also showed that the sequence specific template reactions, metabolic network, and literature parameters were sufficient to predict protein production under different promoters. Recently, aerobic catabolism has been activated in CFPS which increases the usable energy from a carbon sources such as glucose (*1*). The discovery that such complex metabolism could be activated and controlled in CFPS led us to examine the flux distribution of CFPS.

**Figure 2:**
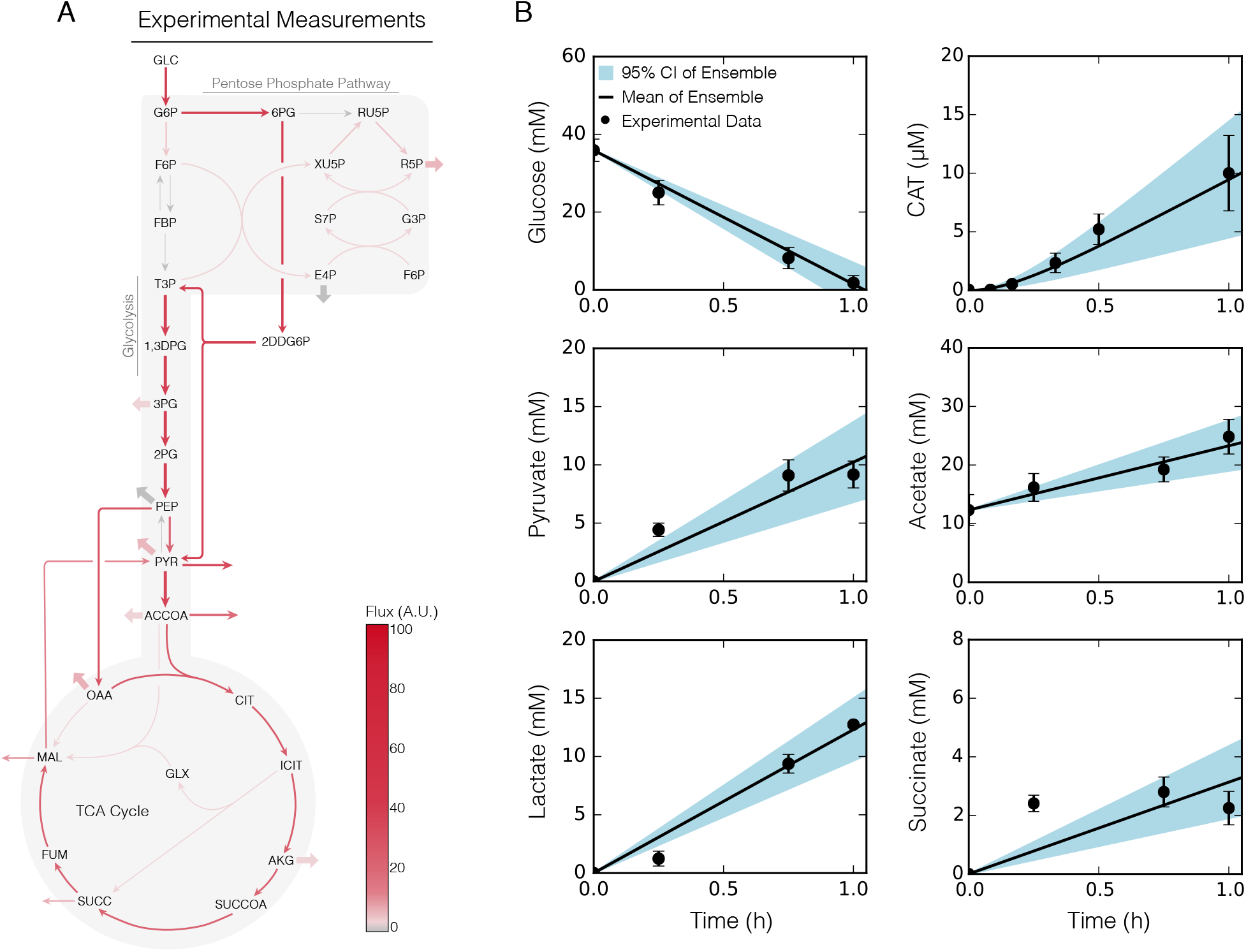
Experimentally constrained simulation of CAT production. CAT was produced under a T7 promoter in CFPS *E. coli* extract for 1 h using glucose as a carbon and energy source. Error bars denote the standard deviation of experimental measurements. The blue region denotes the 95% CI over an ensemble of N = 100 sets, the black line denotes the mean of the ensemble, and dots denote experimental measurements. A. Metabolic flux distribution for CAT production in the presence of experimental constraints for glucose, organic acid and amino acid consumption and production rates. Mean flux across the ensemble, normalized to glucose uptake flux. Thick arrows indicate flux to or from amino acids. B. Central carbon metabolite and CAT measurements versus simulations over a 1 hour time course. The blue region denotes the 95% CI over an ensemble of N = 100 sets, the black line denotes the mean of the ensemble, and dots denote experimental measurements.

### 2.2 Metabolic flux distributions

While there is no cell growth, complex anabolic and catabolic processes still occur during cell free protein synthesis (*36*). To optimize these processes, we must understand the differences in optimal metabolism, and the metabolism occurring in an actual system. Toward this question, we compared the flux distribution of optimal CAT production with experimentally constrained CAT production. The CAT translation rate was optimized without experimental constraints on substrate consumption or byproduct formation to estimate the theoretically optimal metabolic flux distribution. In all cases, the CFPS reaction was supplied with glucose; however, we considered different scenarios for amino acid (AA) supplementation. First, the CFPS reaction was supplied with glucose and amino acids, and was able to synthesize amino acids from glucose (AAs supplied and de novo synthesis). In this case, the flux distribution showed an incomplete TCA cycle, where a combination of glucose and amino acids powered protein expression (Fig. 3A). Glucose was consumed to produce acetyl-coenzyme A, and associated byproducts, while glutamate was converted to alpha-ketoglutarate which traveled to oxaloacetic acid and pyruvate for additional amino acid biosynthesis. Second, the CFPS reaction was supplied with glucose and amino acids, but de novo amino acid biosynthesis was not allowed (AAs supplied w/o de novo synthesis). This scenario was potentially consistent with common cell-free extract preparation protocols which often involve amino acid supplementation; in the presence of supplementation we might expected the enzymes responsible for amino acid biosynthesis to be largely absent from the CFPS reaction. With supplementation and without de novo synthesis, the flux distribution showed no TCA cycle flux with all carbon flux traveling from glucose to acetate. In this case, ATP was produced by a combination of substrate level and oxidative phosphorylation, where ubiquinone was regenerated via either cyo and cyd activity, without relying on succinate dehydrogenase in the TCA cycle (Fig. 3B). These first two cases where amino acids were available had similar performance, and their respective metabolic flux distributions had a 99% correlation. Lastly, the CFPS reaction was supplied with glucose but not amino acids, thus the system was forced to synthesize amino acids de novo from glucose (de novo synthesis only). In the final case, the flux distribution showed a largely complete TCA cycle, and there was diversion of metabolic flux into the Entner-Doudoroff pathway to produce NADPH (Fig. 3C). However, these simulations represent the theoretically optimal metabolic flux distribution, which may not be consistent with what is observed experimentally. Toward this issue, we constrained the feasible solution space with experimental measurements (see Supporting Information) and estimated the optimal CAT metabolic flux distribution.

**Figure 3:**
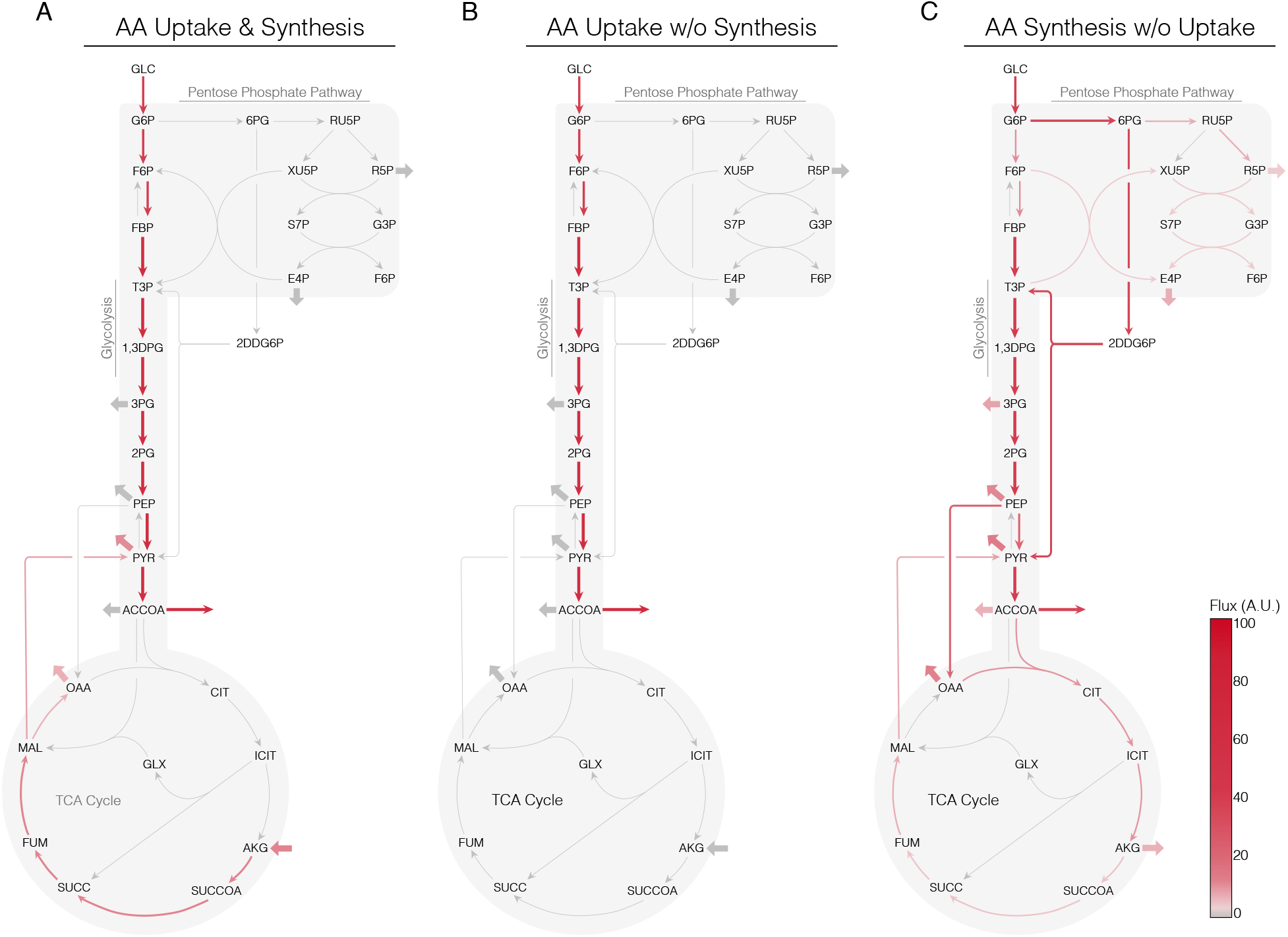
Optimal metabolic flux distribution for CAT production. A. Optimal flux distribution in the presence of amino acid supplementation and de novo synthesis. B. Optimal flux distribution in the presence of amino acid supplementation without de *novo* synthesis. C. Optimal flux distribution with de *novo* amino acid synthesis in the absence of supplementation. Mean flux across the ensemble (N = 100), normalized to glucose uptake flux. Thick arrows indicate flux to or from amino acid biosynthesis pathways.

The experimentally constrained metabolic flux distribution had a 55% correlation with the theoretically optimal flux distribution (Fig. 3). The low similarity suggested several differences between the experimentally constrained and optimal metabolic flux distributions. The largest discrepancy was in oxidative phosphorylation, where the experimental system heavily relied on *cyd* rather than *cyo* to produce ATP through oxidative phosphorylation. The constraint based simulations confirmed that oxidative phosphorylation was active in the cell-free extract, as without it there was no feasible solution within the experimental constraints of the system. The experimentally constrained simulation suggested a high flux through *zwf*, yielding NADPH which was interconverted to NADH via the *pnt1* reaction. This NADH was consumed to convert pyruvate to lactate or to generate ATP via oxidative phosphorylation. In contrast, the optimal solutions with amino acid supplementation had low *zwf* and *pnt1* activity. Surprisingly, folate, purine, and pyrimidine metabolism, along with amino acid biosynthesis, were active in the experimental system, but inactive in the optimal system. In particular, the experimental system had high alanine and glutamine biosynthetic flux (both accumulated in the media), while there was no accumulation of amino acids in the optimal simulations. Lastly, alanine, glutamine, pyruvate, lactate, acetate, malate, and succinate all accumulated in the experimental system, whereas the optimal solution produced (or consumed) only the required amount of metabolites; this accumulation contributed to the difference in the flux distributions. Next, we examined CFPS performance in terms of productivity and energy efficiency.

### 2.3 Analysis of CFPS performance

We analyzed the productivity and energy efficiency for the cell-free production of eight proteins with and without amino acid supplementation (Fig. 4). The expression of each protein was under a P70a promoter, with the exception of CAT which was expressed using a T7 promoter. In all cases, the CFPS reaction was supplied with glucose; however, we considered different scenarios for amino acid (AA) supplementation, similar to the cases considered in the flux distribution: AAs supplied and *de novo* synthesis, AAs supplied w/o *de novo* synthesis, and AA *de novo* synthesis only. Eight proteins, ranging in size, were selected to evaluate CFPS performance: bone morphogenetic protein 10 (BMP10), chloramphenicol acetyltransferase (CAT), caspase 9 (CASP9), dual emission green fluorescent protein (deGFP), prothrombin (FII), coagulation factor X (FX), fibroblast growth factor 21 (FGF21), and single chain variable fragment R4 (scFvR4). An additional case was considered for CAT, where central metabolic fluxes were constrained by experimental measurements of glucose, organic and amino acids (see Supporting Information). Using these model proteins, we developed effective correlation models that predicted the productivity and energy efficiency given the carbon number of the protein. Finally, we independently validated the correlations with a protein not in our original data set: maltose binding protein (MBP).

**Figure 4:**
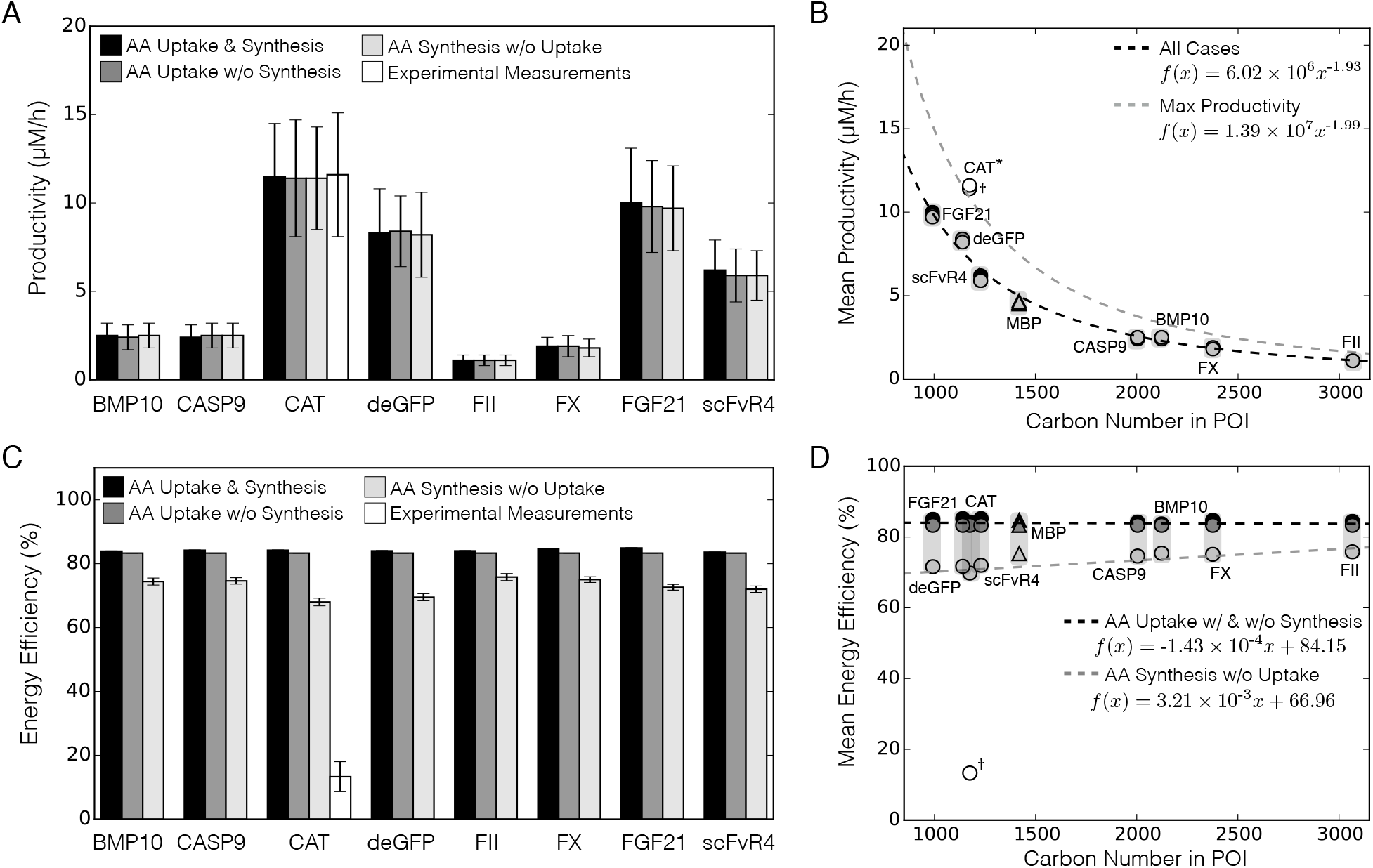
The CFPS performance for eight model proteins with and without amino acid supplementation. A. Mean CFPS productivity for a panel of model proteins with and without amino acid supplementation. B. Mean CFPS productivity versus carbon number for a panel of model proteins with and without amino acid supplementation. Trendline (black dotted line) was calculated across all cases for a P70a promoter (R^2^ = 0.99) and maximum productivity trendline assumed *u* (*κ*) = 1 (grey dotted line; R^2^ = 0.99). C. Mean CFPS energy efficiency for a panel of model proteins with and without amino acid supplementation. D. Mean CFPS energy efficiency versus carbon number for a panel of model proteins with and without amino acid supplementation. Trendline for cases with amino acids (black dotted line) and trendline for without amino acids (grey dotted line; R^2^ = 0.81). Error bars: 95% CI calculated by sampling; asterisk: protein excluded from trendline; dagger: constrained by experimental measurements and excluded from trendline; triangles: first principle prediction and excluded from trendline.

#### 2.3.1 Productivity

The theoretical maximum productivity for proteins expressed using a P70a promoter (*μ*M/h) was inversely proportional to the carbon number (*C_POI_*) and varied between 1 and 12 *μ*M/h for the proteins sampled (Fig. 4A-B). The theoretical maximum productivities, with and without amino acid supplementation, were within a standard deviation of one another for each protein, but varied significantly between proteins. Productivity varied non-linearly with protein length; for instance, BMP10 (424 aa) had a optimal productivity of approximately 2.5 *μ*M/h, whereas the optimal productivity of deGFP (229 aa) was approximately 8.4 *μ*M/h. To examine the influence of protein length, we plotted the mean optimal productivity against the carbon number of each protein (Fig. 4B). The optimal productivity and protein length were related by the power-law relationship *α* × (*C_POI_*)^*β*^, where *α* = 6.02 × 10^6^ *μ*M/(h·carbon number) and *β* = –1.93 for a P70a promoter. Interestingly, CAT did not obey the P70a power-law relationship; the relatively high productivity of CAT was due to its T7 promoter. The higher transcription rate of the T7 promoter increased the steady state level of mRNA by 34%, resulting in a higher productivity. However, CAT expressed under a P70a promoter followed the P70a power-law correlation with a productivity of approximately 8.5 ± 2.3 *μ*M/h (predicted to be 7.2 *μ*M/h by the optimal productivity correlation). Thus, these simulations suggested a promoter specific relationship between the productivity and protein length. However, it was unclear if the productivity correlation was predictive for proteins not considered in the original training set.

We independently validated the productivity correlation by calculating the optimal productivity of MBP (which was not in the original training set) using the full model and the effective correlation model (Fig. 4B). The prediction error was less than 8% for an *a priori* prediction of CFPS productivity using the effective correlation. Thus, the effective productivity correlation could be used as a parameter-free method to estimate optimal productivity for cell-free protein production using a P70a promoter. For CFPS using other promoters, a similar correlation model could be developed. For example, maximal transcription occurs when the promoter model coefficient *u* (*κ*) = 1; the theoretical maximum productivity correlation for maximum promoter activity also followed a power-law distribution (*α* = 1.39 × 10^7^ *μ*M/(h·carbon number) and *β* = –1.99) (Fig. 4B, gray). The CAT value under a T7 promoter was similar to the maximal productivity as *u*_T7_ (*κ*) ≃ 0.91 given the T7 promoter model parameters used in this study (Table 2). Taken together, the maximum optimal productivity of a cell-free reaction was found to be inversely proportional to protein size, following a power-law relationship for proteins expressed under a P70a promoter.

#### 2.3.2 Energy efficiency

The optimal energy efficiency of protein synthesis was independent of protein length, with and without amino acid supplementation (Fig. 4C-D); it was approximately 84% for the model proteins sampled. The relationship was linear, but with negligible slopes: *m_Y_* × (*C_POI_*) + *b_Y_*, where *m_Y_* = –1.43 × 10^−4^ energy efficiency (%)/carbon number for the case with supplementation, and *m_Y_* = 3.21 × 10^−3^ energy efficiency (%)/carbon number for the case without supplementation. The energy efficiency (y-intercept) was calculated at *b_Y_* = 84.15 (%) with supplementation, and *b_Y_* = 66.96 (%) without supplementation. In the presence of amino acids, energy was utilized to power CFPS instead of synthesizing amino acids; thus, a constant energy efficiency was observed regardless of the protein size. In the absence of supplementation, the energy efficiency decreased to between 68% and 76%. In this case, glucose consumption more than doubled (64% increase for CAT) compared to cases supplemented with amino acids; meanwhile, the productivity was similar for each protein (Fig. 4D). Therefore, the energy burden required for synthesizing each amino acid and powering CFPS lowered the energy efficiency. Surprisingly, without amino acid supplementation, proteins with a higher carbon number had marginally higher energy efficiency; however, this linear trend was mostly independent of protein size (R^2^ = 0.82). Lastly, MBP was well predicted by the linear efficiency model with and without amino acid supplementation. The estimated MBP energy efficiency had a maximum error of 6% without supplementation, and an error of 1% in the presence of amino acids.

Experimentally constrained CAT simulations showed suboptimal energy efficiency (Fig. 4D, dagger). CAT production was simulated using the constraint based model in combination with experimental measurements of glucose consumption and organic and amino acid consumption and production rates (Fig. 1B). The experimentally constrained energy efficiency was 16.4 ± 5.6% compared to the theoretical maximum of approximately 84.2 ± 0.1%. Given that the CAT productivity was similar between the simulated and measured systems, differences in the glucose consumption rate and the ATP yield per glucose were likely responsible for the difference between the optimal and experimental systems. The glucose consumption rate was approximately 30 - 40 mM/h in the experimental system (even in the presence of amino acids). On the other hand, the constraint based simulation suggested the optimal glucose consumption rate was significantly less than the observed rate, approximately 1 - 7 mM/h (depending upon amino acid supplementation). In the constraint based simulation, the CFPS reaction produced only acetate as a byproduct, but in the experimental system acetate, lactate, pyruvate, succinate and malate all accumulated during the first hour of production. The energy produced per unit glucose was also different between the optimal and experimentally constrained cases. In the optimal simulation, 12 ATPs were produced per unit glucose (the theoretical maximum for this network was 21), while the experimentally constrained simulation produced only ~4 ATPs per glucose. Thus, approximately 120 - 160 mM ATP/h was produced in the experimental case, in contrast to 12 - 84 mM ATP/h for the optimal case. Thus, the experimental system overproduced ATP. We know from measurements that ATP did not accumulate in the media, which suggested it was consumed by pathways that were not active in the optimal simulation. Thus, CFPS retained an in vivo memory that led to the overconsumption of glucose, and counter intuitively the over production of ATP.

### 2.4 Global sensitivity analysis

We performed global sensitivity analysis to understand which parameters controlled CFPS productivity and energy efficiency (Fig. 5). The translation elongation rate was the most important factor controlling productivity, while RNAP and ribosome abundance had only a modest effect irrespective of amino acid supplementation (Fig. 5A). This suggested that the translation elongation rate, and not transcriptional parameters, controlled productivity. Underwood and coworkers showed that increasing ribosome abundance did not significantly increase protein yields or rates; however, adding elongation factors increased protein synthesis rates by 27% (*37*). In addition, Li et al. increased the productivity of firefly luciferase by 5-fold in PURE CFPS by first improving translation, followed by transcription by adjusting elongation factors, ribosome recycling factor, release factors, chaperones, BSA, and tRNAs (*38*). In examining substrate utilization, glucose consumption was not important for productivity in the presence of amino acid supplementation. However, its importance increased significantly when amino acids were not available. On the other hand, amino acid consumption was only sensitive when *de novo* amino acids biosynthetic reactions were blocked, as these were the only source of amino acids for protein synthesis. The oxygen consumption rate was the most important factor controlling the energy efficiency of cell-free protein synthesis (Fig. 5B). In the model, we assumed that ATP could be produced by both substrate level and oxidative phosphorylation. Jewett and coworkers reported that oxidative phosphorylation still operated in cell-free systems, and that the protein titer decreased from 1.5-fold to 4-fold when oxidative phosphorylation reactions were inhibited in pyruvate-powered CFPS (*1*). Furthermore, we showed that oxidative phosphorylation must be active to simultaneously meet the metabolic and protein production constraints. However, it is unknown how active oxidative phosphorylation is in a glucose-powered cell-free system and its quantitative effect on energy efficiency.

**Figure 5:**
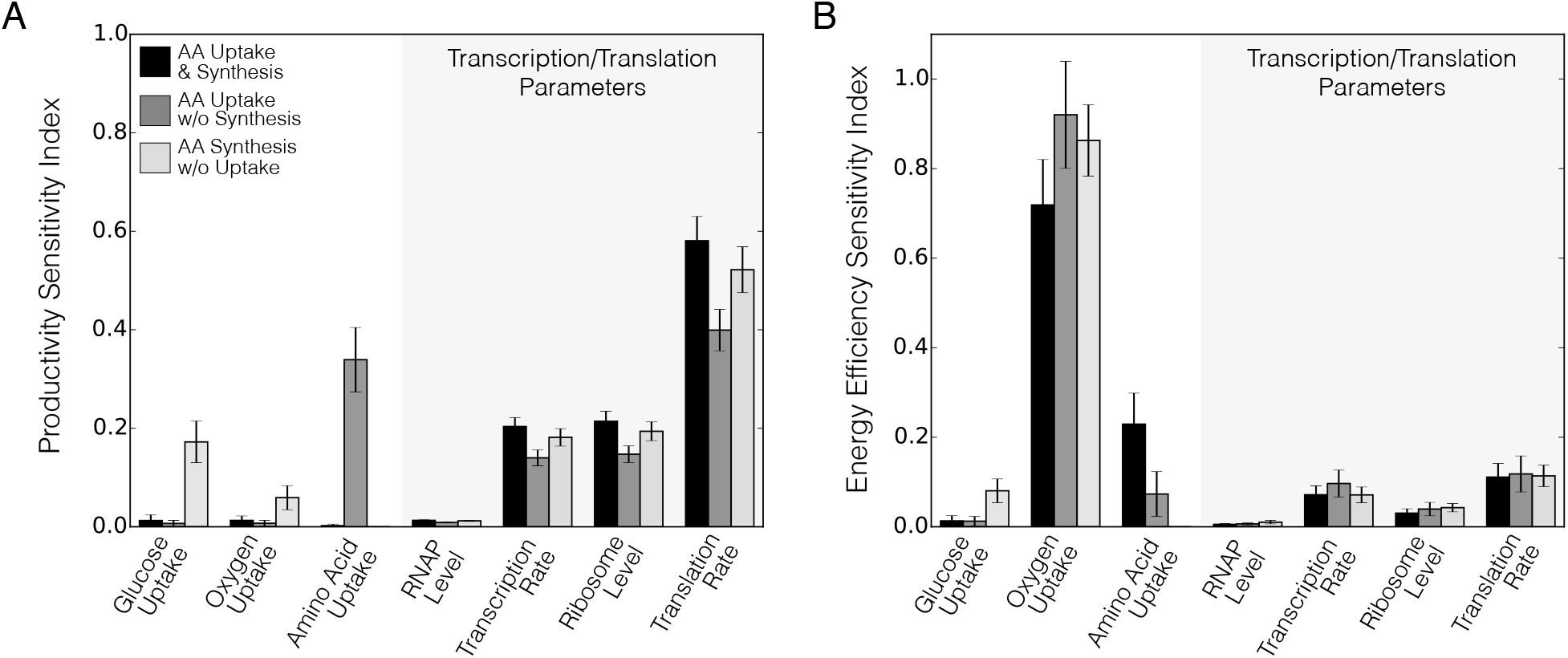
Sensitivity analysis of the cell-free production of CAT. A. Total order sensitivity of the optimal CAT productivity with respect to metabolic and transcription/translation parameters. B. Total order sensitivity of the optimal CAT energy efficiency. Metabolic and transcription/translation parameters were varied for amino acid supplementation and synthesis (black), amino acid supplementation without synthesis (dark grey) and amino acid synthesis without supplementation (light gray). Error bars represent the 95% CI of the total order sensitivity index.

We calculated the optimal CAT energy efficiency as a function of the oxidative phosphorylation flux to investigate the connection between energy efficiency and oxidative phosphorylation (Fig. 6). We calculated energy efficiency across an ensemble of 1000 flux balance solutions by varying the oxygen uptake rate with transcription and translation parameters. Oxidative phosphorylation had a strong effect on the energy efficiency, both with and without amino acid supplementation. In the presence of amino acid supplementation, the energy efficiency ranged from 50% to approximately 84%, depending on the oxidative phosphorylation flux. However, without amino acid supplementation, the energy efficiency dropped to approximately 39%, and reached a maximum of 70%. In the absence of supplementation, a lower energy efficiency was expected for the same oxidative phosphorylation flux, as glucose was utilized for both energy generation and amino acid biosynthesis. In all cases, whenever the energy efficiency was below its theoretical maximum, there was an accumulation of both acetate and lactate. The experimental dataset exhibited a mixture of acetate and lactate accumulation during CAT synthesis, which suggested the CFPS reaction was not operating with optimal oxidative phosphorylation activity. Oxidative phosphorylation is a membrane associated process, while CFPS has no cell membrane. Jewett and coworkers hypothesized that membrane vesicles present in the CFPS reaction carried out oxidative phosphorylation (*1*). Toward this hypothesis, they enhanced the CAT titer by 33% when the reaction was augmented with 10 mM phosphate; they suggested the additional phosphate either enhanced oxidative phosphorylation activity or inhibited phosphatase reactions. The model validated the activity of oxidative phosphorylation in this CFPS system, since without oxidative phosphorylation there was no feasible solution satisfying the constraints of the experimental dataset. However, the number, size, protein loading, and lifetime of these vesicles remains an open area of study.

**Figure 6:**
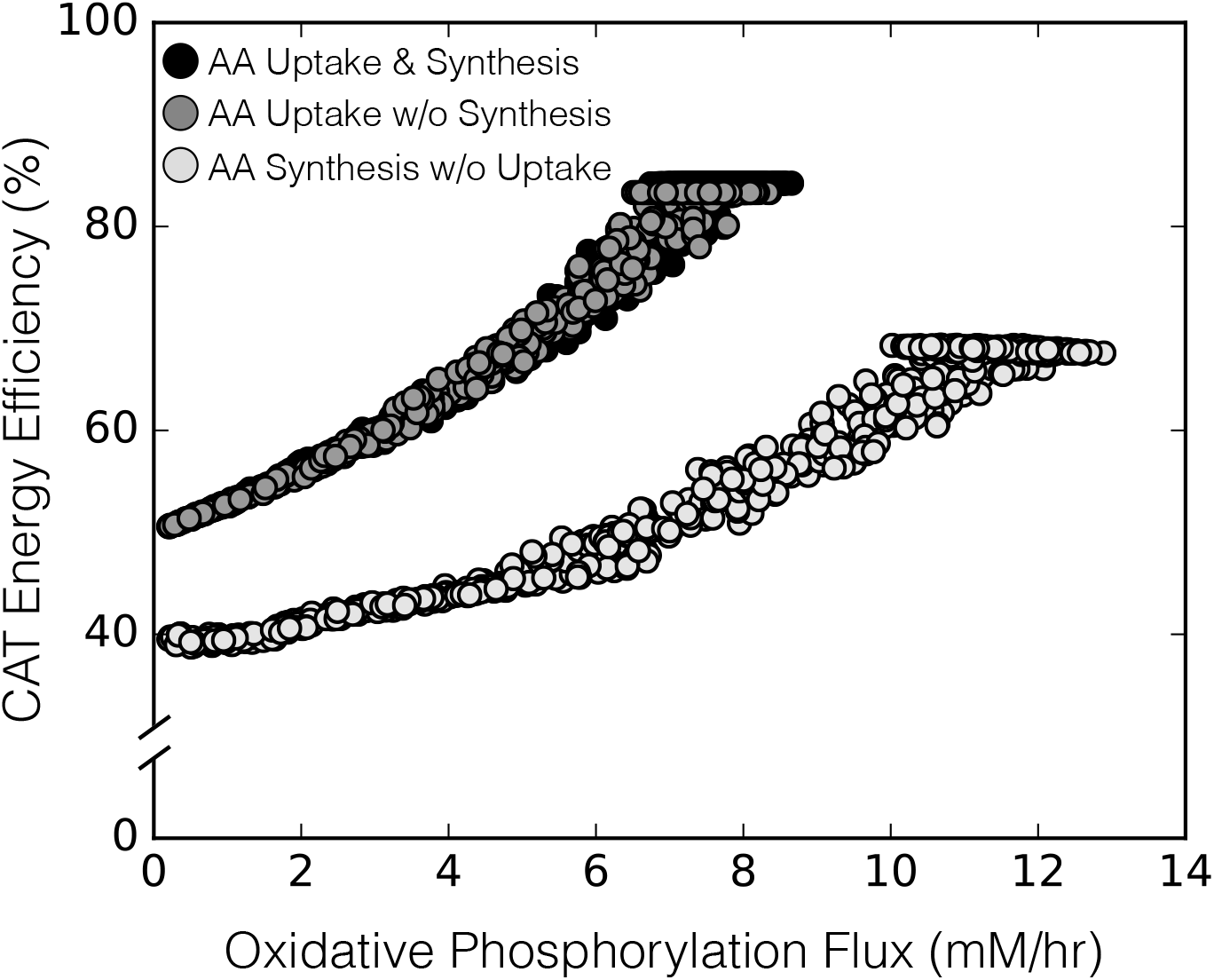
Optimal CAT energy efficiency versus oxidative phosphorylation flux calculated across an ensemble (N = 1000) of flux balance solutions (points). Energy efficiency versus oxidative phosphorylation flux for amino acid supplementation and *de novo* synthesis (black), amino acid supplementation without *de novo* synthesis (dark grey), and de novo amino acid synthesis without supplementation (light gray). The ensemble was generated by randomly varying the oxygen consumption rate from 0.1 to 10 mM/h and randomly sampling the transcription and translation parameters within 10% of their literature values. Each point represents one solution of the model equations.

#### 2.4.1 Potential alternative metabolic optima

Optimal flux distributions predicted using constraint based approaches may not always be unique. Alternative optimal solutions have the same objective value, e.g., productivity, but different metabolic flux distributions. Techniques such as flux variability analysis (FVA) (*39, 40*) or mixed-integer approaches (*41*) can estimate alternative optima. In this study, we used group knockout analysis to estimate potential alternative optimal solutions for CAT production constrained by experimental measurements (Fig. 7). Groups of reactions were removed from the metabolic network, and the translation rate was maximized. The difference between the nominal and altered system was then calculated. Knockout analysis identified pathways required for CAT production; for example, deletion of the glycolysis/gluconeogenesis or oxidative phosphorylation pathways resulted in no CAT production. The absence of CAT production when oxidative phosphorylation was knocked out in the experimentally constrained case confirmed that oxidative phosphorylation was present in the CFPS cell-extract. Likewise, there were pathway knockouts that had no effect on productivity or the metabolic flux distribution, such as removal of isoleucine, leucine, histidine and valine biosynthesis. Globally, the constraint based simulation reached the same optimal CAT productivity for 40% of the pairwise knockouts, while 92% of these solutions had different flux distributions compared with the wild-type. For example, one of the features of the predicted optimal metabolic flux distribution was a high flux through the Entner-Douodoroff (ED) pathway. Removal of the ED pathway had no effect on the CAT productivity compared to the absence of knockouts (Fig. 7A). Pairwise knockouts of the ED pathway and other subgroups (i.e. pentose phosphate pathway, cofactors, folate metabolism, etc.) also resulted in the same optimal CAT productivity. However, there was a difference in the flux distribution with these knockouts (Fig. 7B); thus, alternative optimal metabolic flux distributions exist for CAT production, despite experimental constraints. In addition, knockouts of amino acid biosynthesis reactions had no effect on the productivity with the exception of alanine, aspartate, asparagine, glutamate and glutamine biosynthesis reactions, since amino acids were available in the media. Ultimately, to determine the metabolic flux distribution occurring in CFPS, we need to add additional constraints to the flux estimation calculation. For example, thermodynamic feasibility constraints may result in a better depiction of the flux distribution (*17, 18*), and ^13^C labeling in CFPS could provide significant insight. However, while ^13^C labeling techniques are well established for *in vivo* processes (*42*), application of these techniques to CFPS remains an active area of research.

**Figure 7:**
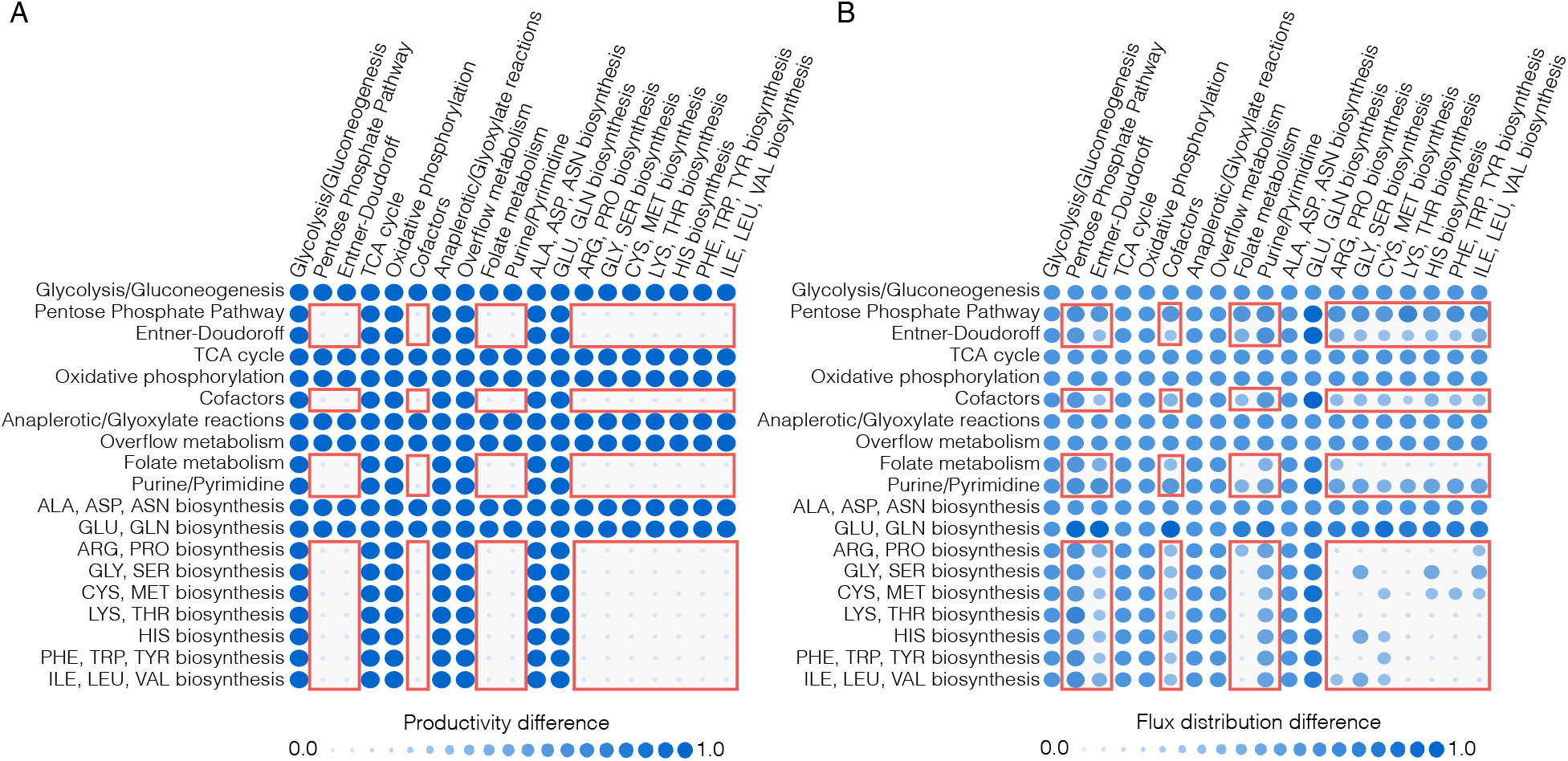
Pairwise knockouts of reaction subgroups in the cell-free network. A. Difference in the CAT productivity in the presence of reaction knockouts compared with no knockouts for experimentally constrained CAT production. B. Difference in the optimal flux distribution in the presence of reaction knockouts compared with no knockouts for experimentally constrained CAT production. The difference between perturbed and wild-type productivity and flux distributions was quantified by the *l*^2^ norm, and then normalized so the maximum change was 1.0. Red boxes indicate potential alternative optimal flux distributions.

### 2.5 Summary and conclusions

In this study, we developed a sequence specific constraint based modeling approach to predict the performance of cell-free protein synthesis reactions. First principle predictions of the cell-free production of CAT and deGFP were in agreement with experimental measurements for two different promoters. While we considered only the P70a and T7 promoters here, we are expanding our library of possible promoters. These promoter models, in combination with the cell-free constraint based approach, could enable the *de novo* design of circuits for optimal functionality and performance. We also developed effective correlation models for the productivity and energy efficiency as a function of protein size that could be used to quickly prototype CFPS reactions. Further, global sensitivity analysis identified the key metabolic processes that controlled CFPS performance; oxidative phosphorylation was vital to energy efficiency, while the translation rate was the most important for productivity. While this first study was promising, there are several issues to consider in future work. First, a more detailed description of transcription and translation reactions has been utilized in genome scale ME models e.g., O’Brien et al (*23*). These template reactions could be adapted to a cell-free system. This would allow us to consider important facets of protein production, such as the role of chaperones in protein folding. We would also like to include post-translation modifications such as glycosylation that are important for the production of therapeutic proteins in the next generation of models. In conclusion, we modeled the cell-free production of a single protein in this study, but sequence specific constraint based modeling could be extended to multi-protein synthetic circuits, RNA circuits or small molecule production.

## Materials and Methods

### Glucose/NMP cell-free protein synthesis

The protein synthesis reaction was conducted using the PANOxSP protocol with slight modifications from that described previously (*43*). The glucose/NMP cell-free protein synthesis reaction was performed using the S30 extract in 1.5-mL Eppendorf tubes (working volume of 15 *μ*L) and incubated in a humidified incubator at 37 ^0^C. The S30 extract was prepared from *E. coli* strain KC6 (A19 ΔtonA ΔtnaA ΔspeA ΔendA ΔsdaA ΔsdaB ΔgshA met+). This K12-derivative has several gene deletions to stabilize amino acid concentrations during the cell-free reaction. The KC6 strain was grown to approximately 3.0 OD595 in a 10-L fermenter (B. Braun, Allentown PA) on defined media with glucose as the carbon source and with the addition of 13 amino acids (alanine, arginine, cysteine, serine, aspartate, glutamate, and glutamine were excluded) (*44*). Crude S30 extract was prepared as described previously (*45*). Plasmid pK7CAT was used as the DNA template for chloramphenical acetyl transferase (CAT) expression by placing the *cat* gene between the T7 promoter and the T7 terminator (*46*). The plasmid was isolated and purified using a Plasmid Maxi Kit (Qiagen, Valencia CA).

All reagents were purchased from Sigma (St. Louis, MO), unless otherwise noted. The initial mixture included 1.2 mM ATP; 0.85 mM each of GTP, UTP, and CTP; 30 mM phosphoenolpyruvate (Roche, Indianapolis IN); 130 mM potassium glutamate; 10 mM ammonium glutamate; 16 mM magnesium glutamate; 50 mM HEPES-KOH buffer (pH 7.5); 1.5 mM spermidine; 1.0 mM putrescine; 34 *μ*g/mL folinic acid; 170.6 *μ*g/mL *E. coli* tRNA mixture (Roche, Indianapolis IN); 13.3 *μ*g/mL pK7CAT plasmid; 100 *μ*g/mL T7 RNA polymerase; 20 unlabeled amino acids at 2-3 mM each; 5 *μ*M l-[U-^14^C]-leucine (Amersham Pharmacia, Uppsala Sweden); 0.33 mM nicotinamide adenine dinucleotide (NAD); 0.26 mM coenzyme A (CoA); 2.7 mM sodium oxalate; and 0.24 volumes of *E. coli* S30 extract. This reaction was modified for the energy source used such that glucose reactions have 30-40 mM glucose in place of PEP. Sodium oxalate was not added since it has a detrimental effect on protein synthesis and ATP concentrations when using glucose or other early glycolytic intermediate energy sources (*47*). The HEPES buffer (pKa ~ 7.5) was replaced with Bis-Tris (pKa ~ 6.5). In addition, the magnesium glutamate concentration was reduced to 8 mM for the glucose reaction since a lower magnesium optimum was found when using a nonphosphorylated energy source (*43*). Finally, 10 mM phosphate was added in the form of potassium phosphate dibasic adjusted to pH 7.2 with acetic acid.

### Protein product and metabolite measurements

Cell-free reaction samples were quenched at specific timepoints with equal volumes of ice-cold 150 mM sulfuric acid to precipitate proteins. Protein synthesis of CAT was determined from the total amount of ^14^C-leucine-labeled product by trichloroacetic acid precipitation followed by scintillation counting as described previously (*35*). Samples were centrifuged for 10 min at 12,000g and 4°C. The supernatant was collected for high performance liquid chromatography (HPLC) analysis. HPLC analysis (Agilent 1100 HPLC, Palo Alto CA) was used to separate nucleotides and organic acids, including glucose. Compounds were identified and quantified by comparison to known standards for retention time and UV absorbance (260 nm for nucleotides and 210 nm for organic acids) as described previously (*35*). The standard compounds quantified with a refractive index detector included inorganic phosphate, glucose, and acetate. Pyruvate, malate, succinate, and lactate were quantified with the UV detector. The stability of the amino acids in the cell extract was determined using a Dionex Amino Acid Analysis (AAA) HPLC System (Sunnyvale, CA) that separates amino acids by gradient anion exchange (AminoPac PA10 column). Compounds were identified with pulsed amperometric electrochemical detection and by comparison to known standards.

### Formulation and solution of the model equations

The sequence specific flux balance analysis problem was formulated as a linear program:

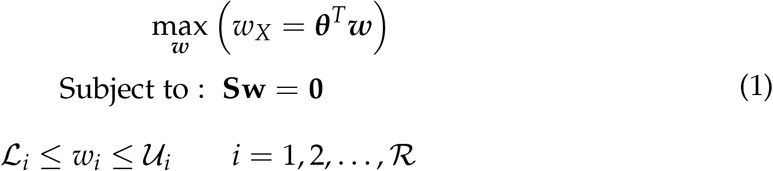

where **S** denotes the stoichiometric matrix 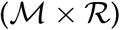, **w** denotes the unknown flux vector 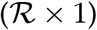, ***θ*** denotes the objective vector 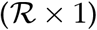 and 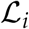 and 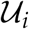 denote the lower and upper bounds on flux *w_i_*, respectively (both 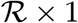 column vectors). Unless otherwise specified, 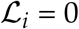 and 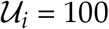 mM/hr. The transcription (T) and translation (X) stoichiometry was modeled using the template reactions of Allen and Palsson (*28*) (Table 1). The objective of the cell free flux balance calculation was to maximize the rate of protein translation, *w_X_*. The total glucose uptake rate was bounded by [0,40 mM/h] according to experimental data, while the amino acid uptake rates were bounded by [0,30 mM/h], but did not reach the maximum flux. Gene and protein sequences were taken from literature and are available in the Supporting Information. The sequence specific flux balance linear program was solved using the GNU Linear Programming Kit (GLPK) v4.55 (*48*). For all cases, amino acid degradation reactions were blocked as these enzymes were likely inactivated during the cell-free extract preparation (*11, 35*). In the absence of *de novo* amino acid synthesis, all amino acid synthesis reactions were set to 0 mM/h. In the experimentally constrained simulations, *E. coli* was grown in the presence of 13 amino acids (alanine, arginine, cysteine, serine, aspartate, glutamate, and glutamine were excluded) (*44*), thus the synthesis reactions responsible for those 13 amino acids were set to 0 mM/h. Lastly, reactions that were knocked out in the host strain used to prepare the extract were removed from the network (ΔspeA, ΔtnaA, ΔsdaA, ΔsdaB, ΔgshA, ΔtonA, ΔendA).

The bounds on the transcription rate 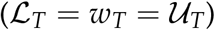 were modeled as:

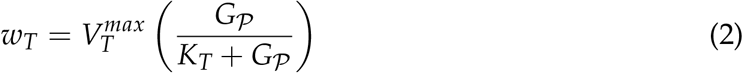

where 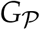 denotes the concentration of the gene encoding the protein of interest, and *K_T_* denotes a transcription saturation coefficient. The maximum transcription rate 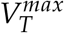 was formulated as:

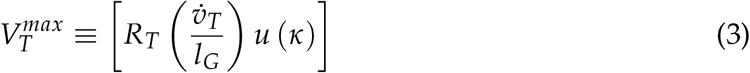

where *R_T_* denotes the RNA polymerase concentration (nM), 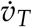 denotes the RNA polymerase elongation rate (nt/h), *I_G_* denotes the gene length (nt). The term *u* (*κ*) (dimensionless, 0 ≤ *u* (*κ*) ≤ 1) is an effective model of promoter activity, where *κ* denotes promoter specific parameters. The general form for the promoter models was taken from Moon *et al*. (*33*); which was based on earlier studies from Bintu and coworkers (*49*), and similar to the genetically structured modeling approach of Lee and Bailey (*50*). In this study, we considered two promoters: T7 and P70a. The promoter function for T7, *u_T7_*, was given by:

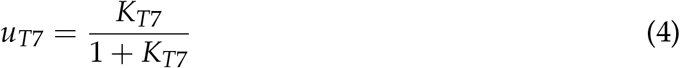

where *K_T7_* denotes a T7 RNA polymerase binding constant. The P70a promoter function *u_P70a_* (which was used for all other proteins) was formulated as:

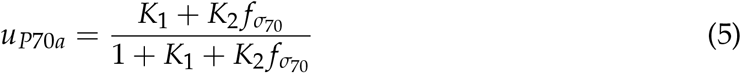

where *K*_1_ denotes the weight of RNA polymerase binding alone, *K*_2_ denotes the weight of RNAP-*σ*_70_ bound to the promoter, and *f*_*p*70_ denotes the fraction of the *σ*_70_ transcription factor bound to RNAP, modeled as a Hill function:

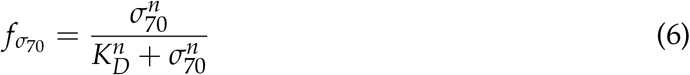

where *σ*_70_ denotes the sigma-factor 70 concentration, *K_D_* denotes the dissociation constant, and *n* denotes a cooperativity coefficient. The values for all promoter parameters are given in Table 2.

The translation rate (*w_X_*) was bounded by:

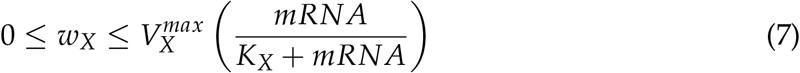

where mRNA^*^ denotes the steady state mRNA abundance and *K_X_* denotes a translation saturation constant. The maximum translation rate 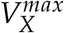 was formulated as:

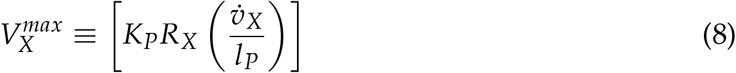

The term *K_P_* denotes the polysome amplification constant, 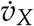 denotes the ribosome elongation rate (amino acids per hour), and *l_P_* denotes the number of amino acids in the protein of interest. The mRNA abundance mRNA was estimated as:

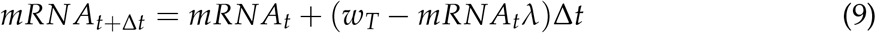

where *λ* denotes the mRNA degradation rate (*h*^−1^). All translation parameters are given in Table 2.

### Calculation of energy efficiency

Energy efficiency 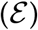 was calculated as the ratio of transcription and translation (weighted by the appropriate energy species coefficients) to ATP generation:

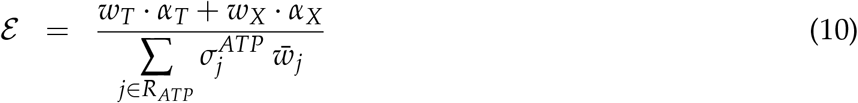

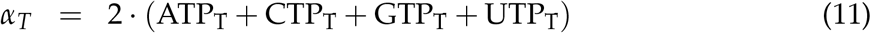

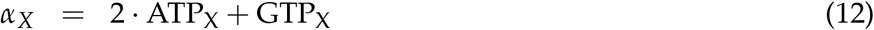

where *α_T_* denotes the energy cost of transcription, *α_X_* denotes the energy cost of translation, *R_ATP_* denotes the set of ATP-producing reactions, and 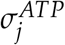 denotes the ATP coefficient for reaction *j*. ATP_T_, CTP_T_, GTP_T_, and UTP_T_ denote the stoichiometric coefficients of each energy species for the transcription of the protein of interest, ATP_X_ and GTP_X_ denote the stoichiometric coefficients of ATP and GTP for the translation of the protein of interest. During transcription and tRNA charging, triphosphate molecules are consumed with monophosphates as byproducts; this is the reason for the factors of 2 on ATP_T_, CTP_T_, GTP_T_, UTP_T_, and ATP_X_

### Quantification of uncertainty

Experimental factors taken from literature, for example macromolecular concentrations or elongation rates, are uncertain. To quantify the influence of this uncertainty on model performance, we randomly sampled the expected physiological ranges for these parameters as determined from literature. An ensemble of flux distributions was calculated for the three different cases we considered: control (with amino acid synthesis and uptake), amino acid uptake without synthesis, and amino acid synthesis without uptake. The flux ensemble was calculated by randomly sampling the maximum glucose consumption rate within a range of 0 to 30 mM/h (determined from experimental data) and randomly sampling RNA polymerase levels, ribosome levels, and elongation rates in a physiological range determined from literature. P70 RNA polymerase levels were sampled between 60 and 80 nM, T70 RNA polymerase levels were sampled between 990 and 1010 nM, ribosome levels between 1.2 and 1.8 μM, the RNA polymerase elongation rate between 20 and 30 nt/s, and the ribosome elongation rate between 1.5 and 3 aa/s (*11, 37*). We generated uniform random samples between an upper (*u*) and lower (*l*) parameter bound of the form:

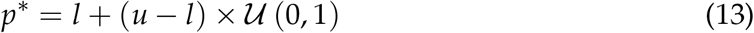

### Global sensitivity analysis

We conducted a global sensitivity analysis using the variance-based method of Sobol to estimate which parameters controlled the performance of the cell-free protein synthesis reaction (*51*). We computed the total sensitivity index of each parameter relative to two performance objectives: productivity of the protein of interest and energy efficiency. We established the sampling bounds for each parameter from literature. We used the sampling method of Saltelli *et al*. (*52*) to compute a family of *N* (2*d* + 2) parameter sets which obeyed our parameter ranges, where *N* was a parameter proportional to the desired number of model evaluations and *d* was the number of parameters in the model. In our case, *N* = 1000 and *d* = 7, so the total sensitivity indices were computed from 16,000 model evaluations. The variance-based sensitivity analysis was conducted using the SALib module encoded in the Python programming language (*53*).

### Potential alternative optimal metabolic flux solutions

We identified potential alternative optimal flux distributions by performing single and pairwise reaction group knockout simulations. Reaction group knockouts were simulated by setting the flux bounds for all the reactions involved in a group to zero and then maximizing the translation rate. We grouped reactions in the cell-free network into 19 subgroups (available in Supporting Information). We computed the difference (*l*^2^-norm) for CAT productivity in the presence and absence of pairwise reaction knockouts. Simultaneously, we computed the difference in the flux distribution (*l*^2^-norm) for each pairwise reaction knockout compared to the flux distribution with no knockouts. Those solutions with the same or similar productivity but large changes in the metabolic flux distribution represent alternative optimal solutions.

## Acknowledgement

This study was supported by the National Science Foundation (MCB-1411715) and the National Science Foundation Graduate Research Fellowship (DGE-1333468) to N.H. This study was also supported by an award from the US Army and Systems Biology of Trauma Induced Coagulopathy (W911NF-10-1-0376) to J.V. for the support of M.V. Lastly, this work was also supported by the Center on the Physics of Cancer Metabolism through Award Number 1U54CA210184-01 from the National Cancer Institute. The content is solely the responsibility of the authors and does not necessarily represent the official views of the National Cancer Institute or the National Institutes of Health.

